# Quantitative genetic basis of floral design in a natural plant population

**DOI:** 10.1101/2020.11.06.371831

**Authors:** Juannan Zhou, Charles B. Fenster, Richard J. Reynolds

## Abstract

The amount of genetic variation of floral traits and the degree to which they are genetically correlated are important parameters for the study of plant evolution. Estimates of these parameters can reveal the effect of historical selection relative to neutral processes such as mutation and drift, and allow us to predict the short-term evolutionary trajectory of a population under various selective regimes. Here, we assess the heritability and genetic correlation of the floral design of a native N. American tetraploid plant, *Silene stellata* (Caryophyllaceae), in a natural population. Specifically, we use a linear mixed model to estimate the genetic parameters based on a genealogy reconstructed from highly variable molecular markers. Overall, we found significant heritabilities in five out of nine studied traits. The level of heritability was intermediate (0.027 – 0.441). Interestingly, the floral trait showing the highest level of genetic variation was previously shown to be under strong sexually conflicting selection, which may serve as a mechanism for maintaining the observed genetic variation. Additionally, we also found prevalent positive genetic correlations between floral traits. Our results suggest that *S. stellata* is capable of responding to phenotypic selection on its floral design, while the abundant positive genetic correlations could also constrain the evolutionary trajectories to certain directions. Furthermore, this study demonstrates the utility and feasibility of marker-based approaches for estimating genetic parameters in natural populations.

## Introduction

A major goal in evolutionary biology is to predict the phenotypic changes of populations under the force of selection. This requires, along with the estimate of selection pressures, detailed understanding of the underlying genetic variation of quantitative traits as well as genetic correlations between traits, as illustrated by the multivariate breeder’s equation (Lande and Arnold, 1983). Furthermore, the pattern of existing genetic variation also reveals the efficacy of historical selection relative to genetic drift, mutation and migration (Lynch and Walsh, 1998).

Since the application of the breeder’s equation to evolutionary studies (Lande and Arnold 1983), extensive efforts have been dedicated to studying the levels of genetic variation in morphological and life history characters in plants (e.g., Ashman and Majetic 2006). Because floral diversity is believed to reflect, in part, pollinator mediated selection(Fenster *et al.*, 2004), and mating system evolution is linked to floral evolution (Barrett, 1998), documentation of within population genetic variation of floral traits has been a historic focus. A survey by Ashman and Majetic (2006) based on studies conducted mostly in controlled environments found an average *h*^*2*^ of 0.39. Although it seems that there is considerable standing genetic variation in floral traits to allow evolutionary response to pollinator-mediated selection (Ashman and Majetic, 2006) and other forces such as herbivores (Strauss and Whittall, 2006), genetic parameters estimated in controlled environments should be interpreted with caution, since plants in general show large phenotypic plasticity (Primack and Kang, 1989), and estimation of heritability in controlled environments could be potentially biased upward due to the lower environmental effects, as well as G×E interactions (Coyne and Beecham, 1987; Conner *et al.*, 2003).

Traditional methods of heritability estimation rely on the covariation of breeding values between close relatives and usually require construction of experimental populations with uniform family structures, e.g. parent-offspring, full-sib, and half-sib designs (Falconer and Mackay, 1996). A large sample size is also required to ensure statistical power (Mitchell-Olds and Rutledge, 1986). Therefore, the cost and labor of this type of breeding program pose significant obstacles to the study of genetic variation in wild plant populations.

Recently, a restricted maximum likelihood (REML) approach termed the “animal model” has been applied to investigate genetic bases of complex traits in wild populations (Kruuk, 2004; Wilson *et al.*, 2010). The animal model is more flexible in that it does not require a fixed family structure in the study population and is able to take into account all relationships in any pedigree. The animal model is therefore expected to provide more statistical power than close-kin comparisons (Kruuk, 2004). It is also less susceptible to confounding effects such as inbreeding, selection and shared environment (Kruuk and Hadfield, 2007), and is more robust to unbalanced datasets than parent-offspring regression approaches (Lynch *et al.*, 1998; Kesson *et al.*, 2008). Given all these advantages, one limitation of the animal model in its application to field studies is the requirement of a fully known pedigree, which is generally very difficult to acquire in natural populations, especially for organisms with complex life history and mating systems. Therefore, despite its success in animal and plant breeding, application of the animal model to evolutionary studies has been relatively rare and mostly has been restricted to bird and large mammal populations that are either under long-term surveillance, or shows behaviors that allow the inference of relatedness between individuals (Milner et al. 2000, Kruuk 2004, Kesson et al. 2008, Akesson et al. 2008). In contrast, the availability of highly variable molecular marker and various statistical methods now makes it possible to implement the animal model with genealogies reconstructed based on individual genotypes (Frentiu *et al.*, 2008; Blonk *et al.*, 2010; Frère *et al.*, 2010). This approach has the potential to expand the range of application of the animal model to wild populations and has been shown to perform well in several animal systems (Frentiu *et al.*, 2008; Blonk *et al.*, 2010; Frère *et al.*, 2010).

In this paper, we focus on a hermaphroditic plant, *Silene stellata*, native to eastern North America. *S. stellata* is an emerging model organism for studying pollinator-plant interaction and the evolution of mutualism (Kephart *et al.*, 2006; Bernasconi *et al.*, 2009) due to its complex interaction with the pollinator/seed predator moth *Hadena ectypa*. A previous study has shown prevalent phenotypic selection on the *S. stellata* floral traits (Zhou *et al.*, 2020). Most notably, the petal dimensions of *S. stellata* were found to be under strong conflicting selection through the two sexual functions of the hermaphroditic plant, mediated by the positive and negative interaction with *H. ectypa*.

Here, we determine the genetic variation as well as the genetic correlation of the *S. stellata* floral design in a natural population of *S. stellata*, based on a genealogy reconstructed using 8 highly variable microsatellite markers. We aim to examine the genetic basis of the *S. stellata* floral design in light of the previously observed phenotypic selection, and to answer three main questions about the evolutionary biology of *S. stellata*: (1) Is the genetic variation consistent with the pattern of selection observed previously? In particular, do the traits under conflicting selection show elevated levels of genetic variation? (2) Is the study *S. stellata* population able to evolve under directional selection? (3) Do the genetic correlation between traits restrict the evolutionary trajectory dictated by the previously observed multivariate phenotypic selection?

## METHODS

### Study population

We focus on a natural population of *Silene stellata* near the University of Virginia’s Mountain Lake Biological Station in Giles County, Virginia, U.S.A. (37.3471, –80.5426, elevation ≈ 1,100–1,300 meters). *Silene stellata* L. is an infrequent, iteroparous, long-lived perennial herb that is distributed throughout the eastern half of the United States. In our Virginia study sites. *S. stellata* flowering occurs from early July through early September and is characterized by panicle inflorescences with white, hermaphroditic, protandrous flowers (average of 25±2.5 ovules/pistil, Reynolds et al. 2009). An individual plant at our study site usually produces multiple stems and on average produces ~40 flowers each flowering season (Reynolds *et al.*, 2012). Outcrossing rate is relatively high based on estimates from both pollinator exclusion experiments (>73%, Reynolds 2008) and mating system estimation using microsatellite markers (~83%, Zhou et al. 2018b).

*S. stellata* exhibits a complex pollination biology, whereby a nocturnal moth, *Hadena ectypa* pollinates the *S. stellata* flowers during its adult stage, but also predates large quantities of *S. stellata* reproductive tissue during its larval stage. The *Silene-Hadena* interaction is further complicated by a guild of generalist pollinator moths, that are equally effective at pollen transfer (Reynolds *et al.*, 2012). In a previous study, we found that the parasitic and mutualistic interaction between *H. ectypa* leads to conflicting selection on the petal dimension through the two sexual functions of *S. stellata* (Zhou *et al.*, 2020). Furthermore, the transition between *H. ectypa* and the copollinating species across the flowering phenology also leads to temporally variable selection.

We measured *S. stellata* plants in four cohorts in 2012 and 2013. In each year, two cohorts of flowering plants were measured in the course of a week during both the early and late flowering season with a one-week separation between experimental periods. We sampled different plants for each cohort and in each year to avoid pseudo-replication, yielding a total number of 227 plants (2012 early, N = 59; 2012 late, N = 58; 2013 early, N = 55; 2014 late, N = 55). All study plants were within a 20-by-20-meter exclosure constructed in a natural *S. stellata* population near Mountain Lake Biological Station (37.348296°, −80.544301°, elevation ≈ 1,100–1,300 m). The two cohorts within a year correspond to periods when *H. ectypa* or the copollinators were the dominant pollinator, respectively (Zhou, Dudash, and Fenster, 2016; Zhou *et al.*, 2020). Our sampling of genotypes to quantify patterns of genetic variation thus take into account potentially different selection pressures that plants in the early and late cohorts experience.

In contrast to the predominant diploidy of *Silene* species distributed in the Old World, the vast majority of N. American *Silene* have been shown to be polyploids, including tetra-, hexa-, and octoploids, with tetraploids being the most common type (Popp and Oxelman, 2007). The N. American polyploid *Silene* species have at least two independent origins. Phylogenetic relationships within the clade containing *S. stellata* are poorly resolved, possibly due to rapid evolution, recombination among homoeologues, and homoplasy (Popp and Oxelman, 2007). The status of (auto- vs. allo-) tetraploidy of *S. stellata* is not well understood. However, the maximum number of alleles amplified per locus was consistently four across all 8 markers in this study and we found no evidence of fixed heterozygotes, supporting a hypothesis of autotetraploidy (Parisod *et al.*, 2010).

### Phenotypic traits

We quantified the pattern of genetic variation and covariation for nine floral characters: (1) corolla tube length, TL; (2) corolla tube width, TW; (3) largest petal length, PL; (4) largest petal width, PW; (5) number of fringes on the distal margin of the petal, FR; (6) anther exsertion, AN; (7) stigma exsertion, ST, (see Figure 1). We measured the distance from the nectary at the base of the flower to the tip of the anther or the stigma and calculated AN and ST as the difference between nectary-anther or nectary-stigma distance and corolla tube length. All floral traits except ST were measured on flowers during the first day of opening (the male phase). Stigma measurements were taken in the first day of the female phase. We counted fringe number (FR) on a randomly chosen petal in which we measured the other six floral traits. Additionally, we recorded the total number of flowers (NF) and floral display height (HT). Measurements were carried out with calipers (0.1 mm) during the course of one week on each plant for PL, PW, FR, TL, TW, ST, AN. Display height was measured as the height of the highest flower to the nearest cm using a meter stick, while NF was quantified by counting the number of flowers open throughout the experimental period. On average five flowers per plant were measured for floral traits (total number of flowers measured, mean number measured per plant: 2012 early, N = 280, 4.75; 2012 late, N = 239, 4.12; 2013 early, N = 294, 5.35; 2014 late, N = 281, 5.11).

**Figure 1.**
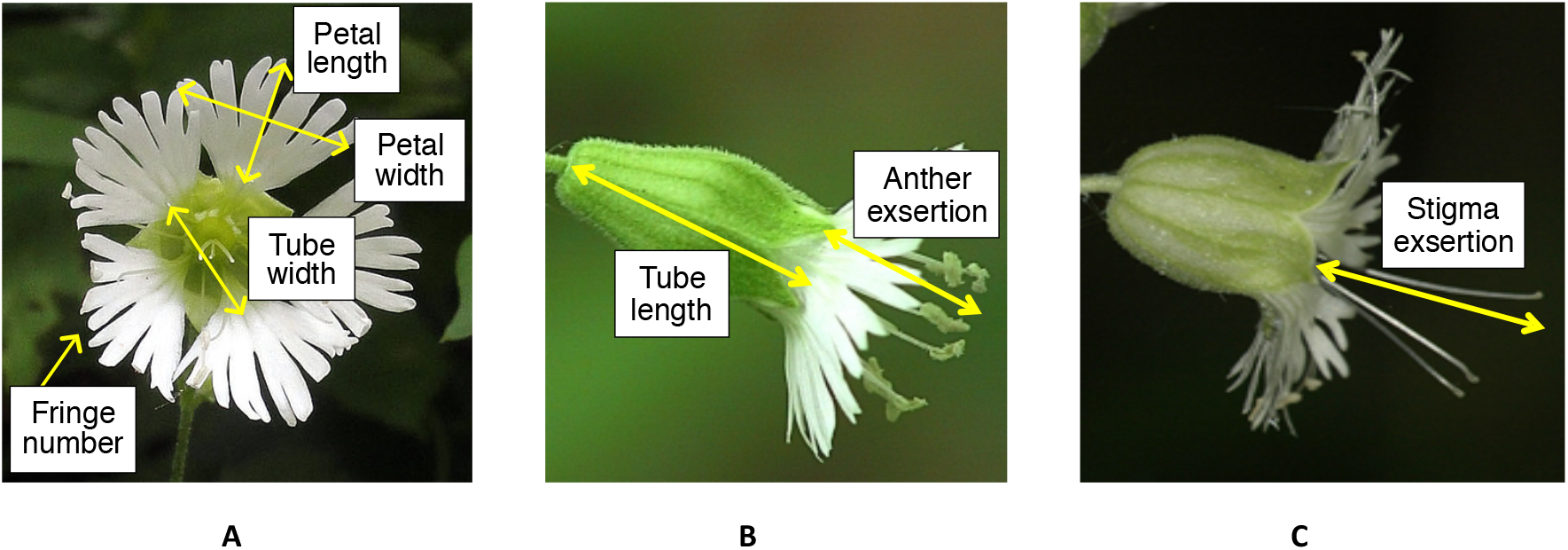
Diagram of *Silene stellata* floral traits measured. All floral traits except stigma exertion were measured on flowers during the first day of opening (the male phase, panel A and B). Stigma exsertion was measured on the first day of the female phase (C).

### Genotyping and genealogy reconstruction

DNA was extracted using the Autogen Plant Kit, following the manufacturer’s protocol. Plants were genotyped on 8 novel microsatellite loci designed for *S. stellata* (Zhou, Dudash, Fenster, *et al.*, 2016). Individual genotypes were visualized and manually scored using GeneMapper V3.7 software (Applied Biosystems). There are two main approaches to replace the known pedigree in the animal model. First, one can construct a matrix, **A**, consisting of relatedness coefficients that are independently estimated for all pairs of individuals. Since the relatedness coefficients are estimated independently for each pair of individuals, compatibility between dyads is not guaranteed (e.g. in examining the relationships between three individuals, A, B and C, the pairs A–B and A–C might be inferred as full-sibs, while the pair B–C might be inferred as half- or non-sib). This incompatibility can lead to the non-positive-definiteness of **A**, resulting in its failure to be inverted for the subsequent REML analysis (Frentiu *et al.*, 2008). Therefore, we implemented the simulated annealing approach using the program COLONY (Jones and Wang, 2010) to reconstruct a genealogy containing all study plants that ensured statistical consistency. COLONY uses the full-pedigree likelihood approach by maximizing the likelihood of the entire pedigree structure rather than individual relationships. Therefore it uses genotype data in a more efficient way and also avoids the troublesome compatibility issue (Jones and Wang, 2010). Additionally, it allows the simultaneous inference of parentage and sibship, which is desirable for our purpose given the extended age structure in the *S. stellata* population (Dudash and Fenster unpublished data). Using COLONY, we generated a genealogy in which all pairs of individuals were classified as full sibs, half sibs, parent-offspring, or unrelated. Since less related pairs of individuals will not be reflected in the estimated genealogy, some information in the data is inherently lost. However, the advantage of genealogy reconstruction is that all pairwise relationships are inherently consistent, and the pedigree data format is compatible with all REML software.

Given that *S. stellata* is a tetraploid, the allelic dosage of partial heterozygotes cannot be readily resolved (for example, an individual showing two alleles A, B on a given locus can have AAAB, AABB or ABBB as the underlying genotype). Additionally, most programs of genealogy reconstruction do not support polyploid data. In the face of these technical difficulties, we transformed the tetraploid genotypes following the protocol described in Wang and Scribner (2014) for application in polyploids. Specifically, we converted the tetraploid genotypes at a codominant microsatellite locus with *k* alleles to diploid phenotypes at *k* dominant “loci”, following the method of Rodzen *et al.* (2004). For example, for a locus with four alleles (A1, A2, A3, A4), an individual with alleles A1, A2, will be coded as {1, 1, 0, 0} at four pseudo-diploid loci, and an individual with alleles A1, A2, A3 will be coded as {1, 1, 1, 0}, and so forth (See Supplemental Table 1 for a detailed example). This approach has been demonstrated to accurately infer sibship, parentage and selfing rate from a typical set of microsatellite markers (Wang and Scribner, 2014).

To assess the reliability of the genealogy reconstructed using the above procedure, we also generated a matrix consisting of independent estimates of pairwise relatedness coefficients, r, using the package POLYRELATEDNESS 1.6 (Huang *et al.*, 2014), designed for autopolyploids. POLYRELATEDNESS uses a maximum likelihood procedure to estimate the vector Δ = [Δ_1_ Δ_2_ Δ_3_ Δ_4_], for every pairwise comparison, where Δ_i_ is the probability that two individuals share *i* alleles that are Identical-By-Descent (IBD). The relatedness coefficient *r* is then calculated as 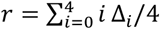. We converted the genealogy generated by COLONY to a relatedness matrix using the statistical package “kinship2” (Therneau and Sinnwell, 2015) in R (R Core Team, 2016). We then compared the two matrices generated by POLYRELATEDNESS and COLONY by running a mantel test in R with 9999 permutations and found a significant correlation (Pearson correlation coefficient *r* = 0.665, *p* < 0.001).

### Mixed model analysis

Univariate variance component analysis was carried out in the REML program Wombat (Meyer, 2007). We fit the following repeated-measure animal model for the seven floral traits using data pooled across years with year and season as fixed effects: *y* = *Xb* + *Z*_*a*_*a* + *Z*_*c*_*c* + *e*, where *y* is the vector of observed phenotypic values, *b* is the vector containing the main effects of year (2012 vs. 2013) and season (early vs. late), *a* is the vector containing individual additive genetic effects, c represents the vector of permanent environmental effects, and *e* contains the residuals. *X*, *Z*_*a*_, and *Z*_*c*_ are design matrices relating appropriate fixed and random effects to individual records. The multiple measurements of a floral trait on a given plant were grouped under the same permanent environmental effect potentially including micro-environmental, maternal, dominance, and epistatic effects on each individual. Permanent environmental effects were not included in the model for number of flowers (NF) and display height (HT), for which only one record was available per plant. The total phenotypic variance *V*_*P*_ is therefore partitioned into additive genetic variance (*V*_*A*_), variance due to permanent environmental effect (*V*_*PE*_), and residual variance (*V*_*E*_). Narrow-sense heritabilities were then calculated using the estimated additive genetic variance as *h*^2^ = *V*_*A*_/(*V*_*A*_ + *V*_*PE*_ + *V*_*E*_). We also reported the repeatability estimates for the seven floral traits as *h*^2^ = (*V*_*A*_ + *V*_*PE*_)/(*V*_*A*_ + *V*_*PE*_ + *V*_*E*_). For comparative purposes, we also report the coefficient of variation (CV), i.e., 100 × additive genetic variance divided by the phenotypic mean.

We specified Wombat to sample 100000 data points from the multivariate normal distribution of ML estimates generated at convergence, to approximate the standard errors for the parameter estimates.

### Genetic correlations

Additive correlations, *r*_*A*_ were estimated using the multivariate version of the animal model described above. The model included only traits for which we observed significant additive genetic components in previous univariate analyses, since *r*_*A*_ is ill-defined if one trait has heritability equal to zero because the genetic correlation between trait *x* and *y* is defined as: 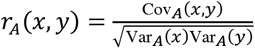, where Cov_*A*_(*x*, *y*)is the genetic covariance (Lynch and Walsh, 1998; Coltman *et al.*, 2005), and. Additionally, we also report phenotypic correlations between pairs of traits based on the outputs of Wombat.

### Correlation of environmental effects between relatives

Since previous work has shown that pollen dispersal distance between plants is as low as 1-2m (Reynolds *et al.*, 2009; Zhou, Zimmer, *et al.*, 2018) and seeds of *S. stellata* mainly disperse passively through gravity, closely related plants could experience common environmental effects due to spatial autocorrelation, which could potentially inflate our genetic component estimates. To rule out this possibility, we compared the relatedness matrix A generated using COLONY with the matrix D which contained pairwise Euclidean distances calculated based on individual GPS coordinates, by running a mantel test with 9999 permutations in R (R Core Team, 2016). No significant correlation was found between the two matrices (Pearson correlation coefficient r = −0.048, *p* > 0.999).

### Statistical tests

We assessed the significance levels of individual parameter estimates as well as comparison of estimates across years by calculating the *z* score, equal to the difference between two estimates divided by the square root of the sum of the squared standard errors. For the purpose of testing significance of single estimates against zero, the formula is reduced to the ratio between the estimate and its standard error. *P*-values of the *z* scores were calculated from a large sample standard normal distribution. We also quantified the association in estimates of *h*^2^, *r*_A_ between years using Pearson’s product-moment correlation (*r*). Additionally, a Mantel test with 9999 permutations was used to compare the two *r*_A_ matrices for 2012 and 2013. All comparisons were performed in R (R Core Team, 2016).

## Results

### Heritability and variance components

We first fit separate REML models for 2012 and 2013. For both years, PW is the only trait with significant heritability (2012: *h*^2^ = 0.530, *p* = 0.06; 2013: *h*^2^ = 0.530, *p* = 0.004). Heritability estimates were similar for all traits across years (Supplemental Table 2; Supplemental Figure 1). Therefore, in the subsequent univariate analyses we pooled the data across years and employed a REML model with year as a fixed effect.

Using the pooled dataset, we found five floral traits (PL, FR, TL, TW, and PW) with significant *h*^2^ (*p* < 0.05), while *h*^2^ of anther exsertion (AN) was marginally significant (*p* = 0.085).. *h*^2^ values ranged from 0 (NF, *p* > 0.99) to 0.441 (PL, *p* < 0.001). Genetic coefficients of variation (CV) ranged between 0.266 (NF) and 14.049. (PL). Repeatabilities were significant in all cases (*p* < 0.05) and ranged from 0.391 (ST) to 0.885 (PL) (Table 1).

**Table 1.** Heritabilities (*h*^*2*^) and repeatabilities (*r*^*2*^) of nine morphological traits of *S. stellata*, with corresponding standard errors (SE), *z* scores (*z*), and significance from zero (*p)*. Parameters were estimated with data pooled across two years including year as a fixed effect, using the univariate animal model. CV_A_: coefficient of variation of additive genetic variance.

| **Trait** | **$h^2$** | **SE $h^2$** | **$z$ $h^2$** | **$p$ $h^2$** | **$r^2$** | **SE $r^2$** | **$z$ $r^2$** | **$p$ $r^2$** | **$CV_A$** |
| --- | --- | --- | --- | --- | --- | --- | --- | --- | --- |
| **PL** | 0.441 | 0.041 | 10.756 | 0.000 | 0.885 | 0.058 | 15.393 | 0.000 | 2.570 |
| **PW** | 0.357 | 0.047 | 7.596 | 0.000 | 0.675 | 0.049 | 13.787 | 0.000 | 2.260 |
| **FR** | 0.360 | 0.047 | 7.660 | 0.000 | 0.720 | 0.051 | 14.073 | 0.000 | 2.312 |
| **TL** | 0.367 | 0.042 | 8.738 | 0.000 | 0.870 | 0.054 | 16.243 | 0.000 | 2.462 |
| **TW** | 0.424 | 0.043 | 9.860 | 0.000 | 0.882 | 0.060 | 14.762 | 0.000 | 2.519 |
| **ST** | 0.027 | 0.131 | 0.206 | 0.837 | 0.391 | 0.136 | 2.867 | 0.004 | 0.453 |
| **AN** | 0.257 | 0.149 | 1.725 | 0.085 | 0.577 | 0.147 | 3.917 | 0.000 | 1.449 |
| **HT** | 0.172 | 0.212 | 0.811 | 0.417 | -<sup>a</sup> | - | - | - | 0.896 |
| **NF** | 0.000 | 0.176 | 0.000 | 1.000 | - | - | - | - | 0.010 |

<sup>a</sup> $r^2$ only available for traits with multiple measurements per plant. PL: petal length; PW: petal width; FR: number of petal fringes; TL: corolla tube length; TW: corolla tube width; ST: stigma exsertion; AN: anther exsertion; HT: display height; NF: number of flowers.

### Genetic correlations

Similar to univariate analyses, we first fit the multivariate animal model for each year independently, then compared the estimates between years. Since no significant differences occurred between years (Supplemental Table 3; Supplemental Figure 2), we pooled the data across years and performed the multivariate animal model using the full model with year as a fixed effect.

Genetic correlations calculated using the pooled dataset were positive in 14 out of 15 comparisons and significant in nine of them (mean *r*_A_ = 0.236 ± 0.295 SE, Table 2). All significant *r*_A_’s were positive and ranged between 0.324 and 0.972. The highest *r*_A_ was TL vs. AN (*r*_A_ = 0.972 ± 0.175 SE, *p* < 0.001), followed by PL vs. PW (*r*_A_ = 0.944 ± 0.260 SE, *p* < 0.001). Petal fringe (FR) was only significantly correlated with PW (*h*^2^ = 0.324 ± 0.161 SE, *p* < 0.05).

**Table 2.** Phenotypic correlations (*r*_*P*_, above the diagonal), additive genetic correlations (*r*_*A*_, below the diagonal), with corresponding standard errors (in parentheses) among six floral traits of *S. stellata*, calculated with data pooled across years using the multivariate animal model with year as a fixed effect. Significance levels were calculated for individual *z* scores based on a large sample standard normal distribution (*** *P*<0.001; ** *P*<0.01; \**P*<0.05; ns, not significant). PL: petal length; PW: petal width; FR: number of petal fringes; TL: corolla tube length; TW: corolla tube width; ST: stigma exsertion; AN: anther exsertion; HT: display height; NF: number of flowers.

|  |  | **Attraction** |  |  | **Pollen transfer** |  |  |
| --- | --- | --- | --- | --- | --- | --- | --- |
|  |  | PL | PW | FR | TL | TW | AN |
| **Attraction** | PL | - | 0.559 (0.047)<br>*** | 0.19 (0.056)<br>*** | 0.388 (0.052)<br>*** | 0.39 (0.049)<br>*** | 0.277 (0.049)<br>*** |
|  | PW | 0.944 (0.260)<br>*** | - | 0.142 (0.054) ** | 0.208 (0.05)<br>*** | 0.527 (0.043)<br>*** | 0.353 (0.045)<br>*** |
|  | FR | 0.310 (0.187)<br>ns | 0.324 (0.161)<br>* | - | 0.08 (0.055)<br>ns | 0.116 (0.054)<br>* | 0.141 (0.051)<br>** |
| **Pollen transfer** | TL | 0.466 (0.198)<br>* | 0.337 (0.145)<br>* | 0.178 (0.145) ns | - | -0.208 (0.053) *** | 0.487 (0.039)<br>*** |
|  | TW | 0.774 (0.257)<br>** | 0.663 (0.177)<br>*** | 0.283 (0.165) ns | -0.062 (0.138) ns | - | 0.157 (0.05)<br>** |
|  | AN | 0.626 (0.204)<br>** | 0.474 (3.12)<br>** | 0.296 (0.160) ns | 0.972 (0.175)<br>*** | 0.163 (0.143)<br>ns | - |

Similar to genetic correlations, values of phenotypic correlations *r*_p_ were positive except for TL vs. TW (*r*_P_ = −0.208 ± 0.053 SE, *p* < 0.001) (Table 2). Mean *r*_P_ = 0.2538 ± 0.202 SD. The only insignificant *r*_P_ was TL vs. FR (*r*_P_ = 0.08 ± 0.055 SE, *p* > 0.05). The average squared genetic correlation (*r*^*2*^_A_) was 0.28, while the average squared phenotypic correlation (*r*^*2*^_P_) was 0.10 (Cheverud 1988). A Mantel test with 999 permutations revealed significant correlation between the genetic and phenotypic correlation matrices (Pearson correlation coefficient *r* = 0.91, *p* < 0.001).

## Discussion

Although statistically more powerful and flexible than traditional methods such as parent-offspring regression, the requirement of a known pedigree has impeded the application of the animal model to natural plant populations. There have been a number of studies that attempted to replace the known pedigree with one reconstructed from molecular genotypes (Frentiu et al. 2008, Blonk et al. 2010, Frère et al. 2010). These results generally support the efficacy of the pedigree-free approach in estimating genetic components. Here we reconstructed a genealogy for 227 *S. stellata* individuals based on 11 highly variable microsatellite loci and observed prevalent genetic variation as well as covariation in the floral traits of *S. stellata*.

One major challenge in using molecular genealogy in an animal model analysis is the statistical error in the inference of pairwise relationship due to the limitation of genetic markers and the genealogy reconstruction algorithm, because the pedigree is assumed by the model to be known without error. Little empirical work has been conducted to examine how erroneous relationship inference affects the animal model. However, we used a microsatellite marker set that was highly polymorphic (mean number of alleles per locus = 22.6). Additionally, we observed a fair range of relatedness including unrelated, half-sib, full-sib and parent-offspring in the reconstructed genealogy. The high resolution of our marker set and the variation in relatedness may help to minimize the effect of erroneous relationships on our inference of genetic parameters (Ritland, 1996; Frentiu *et al.*, 2008).

We observed significant heritabilities, *h*^2^ in five out of nine traits of *S. stellata*. Additionally, anther exsertion (AN) was marginally significant (*h*^2^ = 0.257 ± 0.149 SD, *p* = 0.085). The *h*^2^’s were intermediate (0.027–0.441) for all the remaining floral traits. Among all floral traits, stigma exsertion (ST) had the lowest heritability (*h*^2^ = 0.027 ± 0.131 SD, *p* > 0.10). One likely explanation for this low *h*^2^ other than strong historical selection, is that the protandrous *S. stellata* flowers transition to the female phase after the one-day male-phase through slow extension of the style. Thus, although we made all measurements of ST on flowers during the first day of the female stage, the exact time of measurements and possible asynchrony in growth rates of the styles could introduce additional environmental variation. We did not find significant *h*^2^ in display height (HT) and number of flowers (NF). One likely cause for the lack of genetic components in NF and HT could be that *S. stellata* is a long-lived perennial, and phenotypic variation in these two traits mainly reflects variation in plant age and local microhabitat, rather than the underlying genetic factors.

Our mean estimate of *h*^2^ over all nine traits was 0.27, while mean *h*^2^ of the seven floral traits was 0.32. These estimates were lower than the mean of 0.39 reported by Ashman and Majetic (2006). This is not surprising, as 66% percent of the studies surveyed by Ashman and Majetic (2006) were conducted in controlled environments and high environmental variance in field conditions might contribute to our lower observed *h*^2^. Additionally, in a previous study, we also observed substantial neutral genetic variation in three local *S. stellata* populations (Zhou, Dudash, *et al.*, 2018). Our assessment of genetic variation in morphological traits here seems to be concordant with this previous finding, although there is generally a weak relationship between levels of variation of neutral markers and quantitative traits (Reed *et al.*, 2002).

In a previous study (Zhou *et al.*, 2020), we found significant directional selection on several floral traits of *S. stellata*, including petal length, petal width, and fringe number, as well as vegetative traits including number of flower and display height. Our finding of prevalent heritability in the *S. stellata* floral traits indicate the potential for short-term evolutionary response to contemporary selective pressures. On the other hand, although number of flowers were consistently under the strongest positive selection, the absence of genetic variation of this trait precludes its response to the observed selective pressure.

Importantly, in Zhou *et al.* (2020), we also found conflicting selection pressures exerted on the petal dimensions through the two sexual functions. Remarkably, the floral trait under the strongest conflicting selection (petal length) is also found to have the highest heritability in this paper. Additionally, an earlier multi-year study of the same *S. stellata* population showed that certain floral traits were under phenotypic selection significant only in certain years (Zhou *et al.*, 2020). Together, this indicates that the prevalent additive genetic variation in our study population could have resulted from conflicting selection exerted through different fitness components of the plant and/or the temporal fluctuation of selective pressures. Finally, while G×E interactions could also be responsible for the maintenance of genetic variation (Gillespie and Turelli, 1989), the high similarities in *h*^2^ and *r*_A_ across years do not seem to support this explanation.

We found significant genetic correlations, *r*_A_ in nine out of 15 pairwise comparisons. We found a mean *r*_A_ of 0.45, which is higher than Ashman and Majetic (2006) (Mean±ST = 0.33±0.03). Following Cheverud (1988), we compared the phenotypic and genetic correlation matrices using the average squared correlation coefficients. The average squared genetic correlation was 0.28, while the average squared phenotypic correlation was 0.10. This difference is very close to the mean difference of 0.20 reported by Cheverud (1988), and may be caused by sampling error given the limitation of a pedigree-free animal model. Alternatively, the observed differences could be caused by significant environmental variation in the natural environment (Willis *et al.*, 1991), since the majority of cases studied by Cheverud (1988) were conducted in laboratory or agricultural environments.

The significant *r*_A_’s were uniformly positive. This suggests that the floral design of *S. stellata* could potentially respond harmoniously to natural selection for flower size. On the other hand, divergence of character states through selection may be constrained along axes determined by the abundant genetic correlations (Falconer, 1981). Specifically, genetic correlations between traits associated with pollinator attraction (PL, PW, FR in this study) (Bell, 1985; Ashman and Majetic, 2006; Mitchell *et al.*, 2015) and traits associated with pollen transfer (TL, TW, AN, ST in this study) (Nilsson, 1988; Campbell *et al.*, 1991, 1996; Ashman and Majetic, 2006; Reynolds *et al.*, 2010) were significant in six of nine cases. These positive genetic correlations could hamper independent evolution of different functional modules under potentially opposing selection pressures by various selective agents (e.g., mutualists vs. antagonists) and through the male vs. female function of the hermaphroditic flower (Strauss and Whittall, 2006; Steven *et al.*, 2007). In contrast, the lack of genetic correlation between certain pairs of floral traits could provide additional minor axes for divergence of character states. For example, the *r*_A_ between corolla tube length (TL) and width (TW) is close to zero (*r*_A_ = −0.062 ± 0.138 ST, *p* > 0.10) while both traits had significant *h*^2^. This indicates that these two dimensions of the *S. stellata* corolla tube could potentially respond independently to opposing selection pressures.

In conclusion, our results show the study *S. stellata* population exhibit prevalent moderate genetic variation in its floral design, which can be attributed to the previously observed conflicting and variable selection regime. This shows the study population is capable of responding to directional selection in the face of environmental change, whereas the genetic correlation between traits could also impose evolutionary constraints in certain multivariate selection scenarios.

---

**Supplemental Table 1.** Sample pseudo-diploid input file containing actual genotypes of 10 individuals at locus 3R with 20 alleles.

| ID | Original tetraploid genotype |  |  |  | Transformed genotypes at 20 pseudo-diploid dominant loci |  |  |  |  |  |  |  |  |  |  |  |  |  |  |  |  |  |  |  |
| --- | --- | --- | --- | --- | --- | --- | --- | --- | --- | --- | --- | --- | --- | --- | --- | --- | --- | --- | --- | --- | --- | --- | --- | --- |
|  | Locus 3R (bp) |  |  |  | 3R-142 | 3R-145 | 3R-148 | 3R-151 | 3R-154 | 3R-157 | 3R-160 | 3R-163 | 3R-166 | 3R-169 | 3R-172 | 3R-175 | 3R-178 | 3R-181 | 3R-184 | 3R-187 | 3R-190 | 3R-193 | 3R-196 | 3R-199 |
| 12001 | 148 | 160 | 169 |  | 0 | 0 | 1 | 0 | 0 | 0 | 1 | 0 | 0 | 1 | 0 | 0 | 0 | 0 | 0 | 0 | 0 | 0 | 0 | 0 |
| 12011 | 145 | 148 | 154 | 169 | 0 | 1 | 1 | 0 | 1 | 0 | 0 | 0 | 0 | 1 | 0 | 0 | 0 | 0 | 0 | 0 | 0 | 0 | 0 | 0 |
| 12012 | 148 | 157 | 163 |  | 0 | 0 | 1 | 0 | 0 | 1 | 0 | 1 | 0 | 0 | 0 | 0 | 0 | 0 | 0 | 0 | 0 | 0 | 0 | 0 |
| 12021 | 151 | 154 | 172 | 175 | 0 | 0 | 0 | 1 | 1 | 0 | 0 | 0 | 0 | 0 | 1 | 1 | 0 | 0 | 0 | 0 | 0 | 0 | 0 | 0 |
| 12031 | 145 | 148 | 169 | 181 | 0 | 1 | 1 | 0 | 0 | 0 | 0 | 0 | 0 | 1 | 0 | 0 | 0 | 1 | 0 | 0 | 0 | 0 | 0 | 0 |
| 12041 | 148 | 154 | 187 |  | 0 | 0 | 1 | 0 | 1 | 0 | 0 | 0 | 0 | 0 | 0 | 0 | 0 | 0 | 0 | 1 | 0 | 0 | 0 | 0 |
| 12042 | 148 | 154 | 157 | 160 | 0 | 0 | 1 | 0 | 1 | 1 | 1 | 0 | 0 | 0 | 0 | 0 | 0 | 0 | 0 | 0 | 0 | 0 | 0 | 0 |
| 12101 | 148 | 151 | 166 | 172 | 0 | 0 | 1 | 1 | 0 | 0 | 0 | 0 | 1 | 0 | 1 | 0 | 0 | 0 | 0 | 0 | 0 | 0 | 0 | 0 |
| 12111 | 148 | 157 | 175 |  | 0 | 0 | 1 | 0 | 0 | 1 | 0 | 0 | 0 | 0 | 0 | 1 | 0 | 0 | 0 | 0 | 0 | 0 | 0 | 0 |
| 12121 | 148 | 151 | 163 | 169 | 0 | 0 | 1 | 1 | 0 | 0 | 0 | 1 | 0 | 1 | 0 | 0 | 0 | 0 | 0 | 0 | 0 | 0 | 0 | 0 |

**Supplemental Table 2.** Heritabilities (*h*^*2*^) and corresponding standard errors (SE), *z* scores (*z*), and significance from zero (*p)* of nine morphological traits of *S. stellata* estimated for year 2012 and 2013 based on 11 microsatellite loci, using the univariate animal model. Also shown are the *z* scores and significance levels (*p)* for comparison of *h*^*2*^ across years. Significance levels were calculated for individual *z* scores based on a large sample standard normal distribution. PL: petal length; PW: petal width; FR: number of petal fringes; TL: corolla tube length; TW: corolla tube width; ST: stigma exsertion; AN: anther exsertion; HT: display height; NF: number of flowers.

| Trait | 2012 |  |  |  | 2013 |  |  |  | 2012 vs. 2013 |  |
| --- | --- | --- | --- | --- | --- | --- | --- | --- | --- | --- |
| | $h^2$ | SE | $z$ | $p$ | $h^2$ | SE | $z$ | $p$ | $z$ | $p$ |
| **PL** | 0.366 | 0.599 | 0.611 | 0.541 | 0.339 | 0.319 | 1.063 | 0.288 | 0.040 | 0.968 |
| **PW** | 0.530 | 0.284 | 1.866 | 0.062 | 0.538 | 0.185 | 2.908 | 0.004 | -0.024 | 0.981 |
| **FR** | 0.344 | 0.495 | 0.695 | 0.487 | 0.363 | 0.350 | 1.037 | 0.300 | -0.031 | 0.975 |
| **TL** | 0.351 | 0.427 | 0.822 | 0.411 | 0.310 | 0.289 | 1.073 | 0.283 | 0.080 | 0.937 |
| **TW** | 0.332 | 0.335 | 0.991 | 0.322 | 0.350 | 0.277 | 1.264 | 0.206 | -0.041 | 0.967 |
| **ST** | 0.000 | 0.251 | 0.000 | 1.000 | 0.001 | 0.334 | 0.003 | 0.998 | -0.002 | 0.998 |
| **AN** | 0.437 | 0.385 | 1.135 | 0.256 | 0.411 | 0.270 | 1.522 | 0.128 | 0.055 | 0.956 |
| **HT** | 0.052 | 0.389 | 0.134 | 0.894 | 0.044 | 0.399 | 0.110 | 0.912 | 0.014 | 0.989 |
| **NF** | 0.000 | 0.332 | 0.000 | 1.000 | 0.089 | 0.382 | 0.233 | 0.816 | -0.176 | 0.860 |

**Supplemental Table 3.** Genetic correlations (*r*_*A*_) and corresponding standard errors (SE), *z* scores and significance from zero (*p)* between six morphological traits of *S. stellata* estimated for year 2012 and 2013, using the multivariate animal model. Also shown are the *z* scores and significance levels for comparisons of *r*_*A*_ across years. Significance levels were calculated for individual *z* scores based on a large sample standard normal distribution. PL: petal length; PW: petal width; FR: number of petal fringes; TL: corolla tube length; TW: corolla tube width; AN: anther exsertion.

|  | 2012 |  |  |  | 2013 |  |  |  | 2013 vs. 2013 |  |
| --- | --- | --- | --- | --- | --- | --- | --- | --- | --- | --- |
| | $r_A$ | SE | $z$ | $p$ | $r_A$ | SE | $z$ | $p$ | $z$ | $p$ |
| **PL vs. PW** | 0.933 | 0.762 | 1.224 | 0.221 | 0.915 | 0.417 | 2.194 | 0.028 | 0.021 | 0.983 |
| **PL vs. FR** | 0.406 | 0.483 | 0.841 | 0.401 | 0.354 | 0.314 | 1.127 | 0.260 | 0.090 | 0.928 |
| **PL vs. TL** | 0.340 | 0.448 | 0.759 | 0.448 | 0.350 | 0.354 | 0.989 | 0.323 | -0.018 | 0.986 |
| **PL vs. TW** | 0.852 | 0.788 | 1.081 | 0.280 | 0.860 | 0.509 | 1.690 | 0.091 | -0.009 | 0.993 |
| **PL vs. AN** | 0.525 | 0.451 | 1.164 | 0.244 | 0.551 | 0.357 | 1.543 | 0.123 | -0.045 | 0.964 |
| **PW vs. FR** | 0.380 | 0.354 | 1.073 | 0.283 | 0.312 | 0.237 | 1.316 | 0.188 | 0.160 | 0.873 |
| **PW vs. TL** | 0.365 | 0.343 | 1.064 | 0.287 | 0.308 | 0.233 | 1.322 | 0.186 | 0.137 | 0.891 |
| **PW vs. TW** | 0.657 | 0.387 | 1.698 | 0.090 | 0.665 | 0.280 | 2.375 | 0.018 | -0.017 | 0.987 |
| **PW vs. AN** | 0.417 | 0.254 | 1.642 | 0.101 | 0.410 | 0.210 | 1.952 | 0.051 | 0.021 | 0.983 |
| **FR vs. TL** | 0.158 | 0.293 | 0.539 | 0.590 | 0.186 | 0.257 | 0.724 | 0.469 | -0.072 | 0.943 |
| **FR vs. TW** | 0.386 | 0.421 | 0.917 | 0.359 | 0.338 | 0.286 | 1.182 | 0.237 | 0.094 | 0.925 |
| **FR vs. AN** | 0.319 | 0.337 | 0.947 | 0.344 | 0.267 | 0.225 | 1.187 | 0.235 | 0.128 | 0.898 |
| **TL vs. TW** | -0.091 | 0.266 | -0.342 | 0.732 | -0.069 | 0.229 | -0.301 | 0.763 | -0.063 | 0.950 |
| **TL vs. AN** | 0.916 | 0.606 | 1.512 | 0.131 | 0.949 | 0.486 | 1.953 | 0.051 | -0.042 | 0.966 |
| **TW vs. AN** | 0.238 | 0.267 | 0.891 | 0.373 | 0.222 | 0.215 | 1.033 | 0.302 | 0.047 | 0.963 |

**Supplemental Figure 1.**
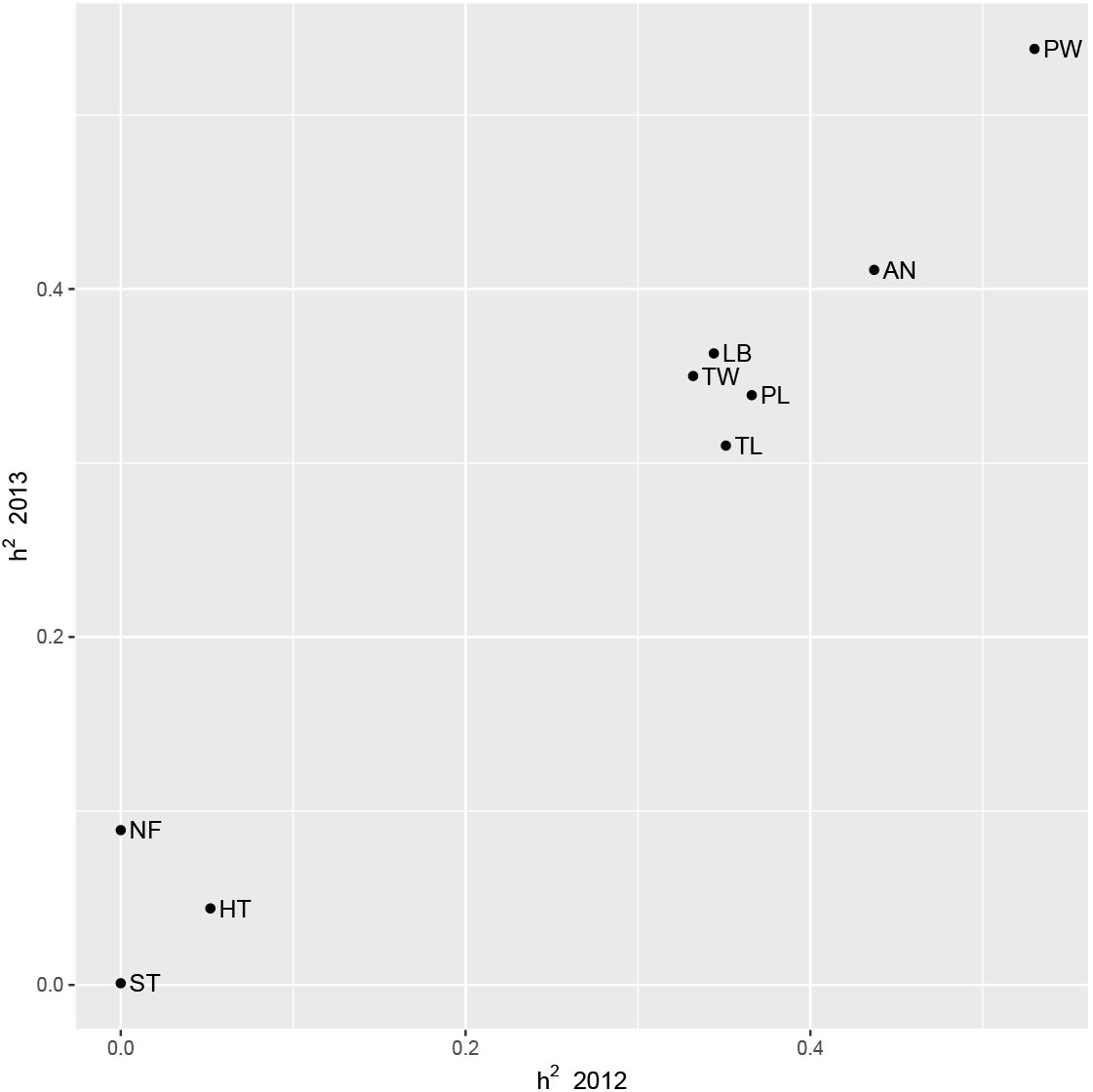
Association between heritabilities (*h*^*2*^) estimated for year 2012 and 2013, using the univariate animal model. Pearson correlation coefficient r = 0.879, *p* < 0.01; Spearman rank correlation ρ= 0.854, *p* < 0.01. PL: petal length; PW: petal width; FR: number of petal fringes; TL: corolla tube length; TW: corolla tube width; ST: stigma exsertion; AN: anther exsertion; HT: display height; NF: number of flowers.

**Supplemental Figure 2.**
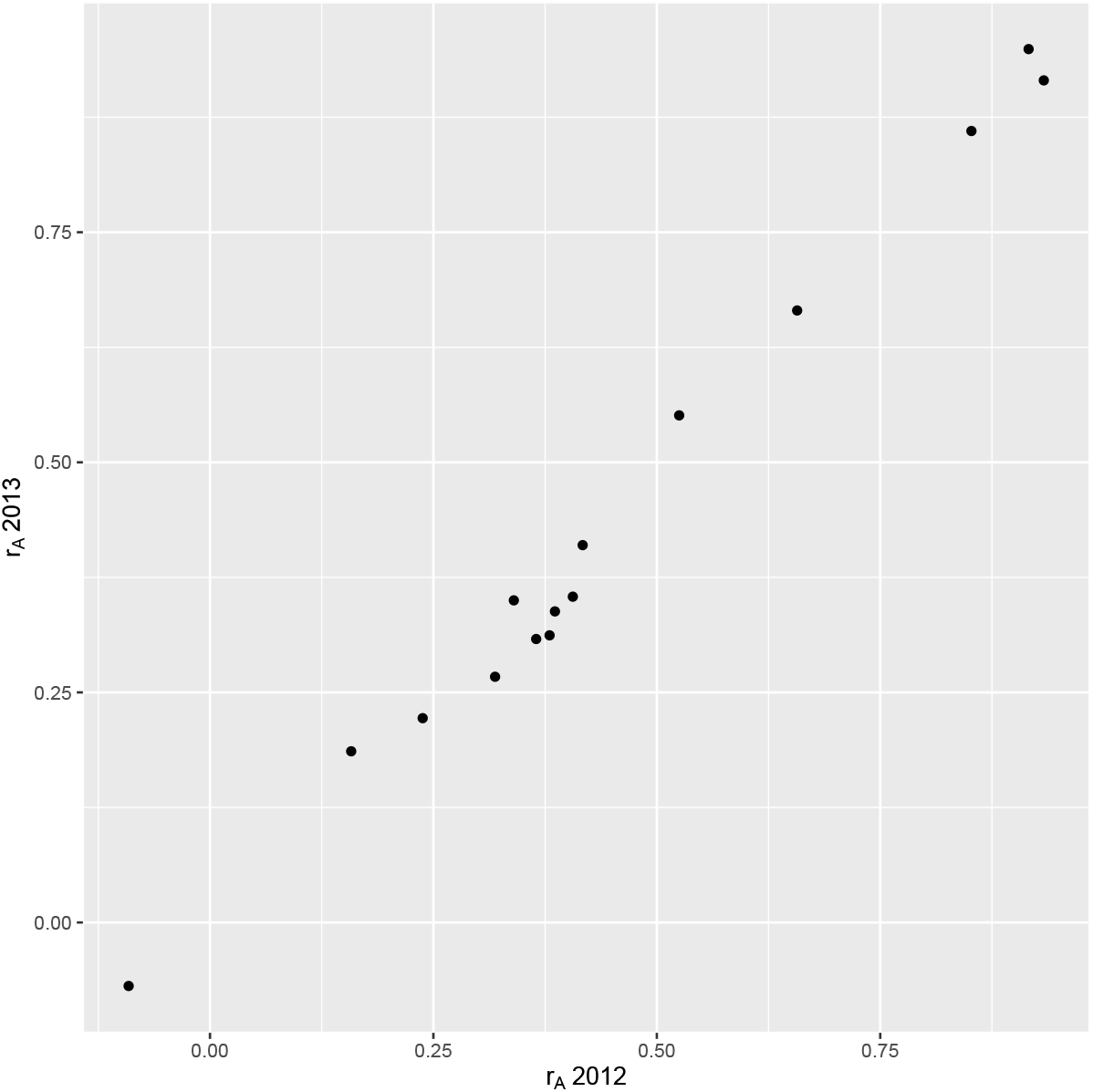
Association between genetic correlations (*r*_*A*_) estimated for year 2012 and 2013 using the multivariate animal model. Pearson correlation coefficient r = 0.978, *p* < 0.001; Spearman rank correlation ϱ = 0.975, *p* < 0.001. PL: petal length; PW: petal width; FR: number of petal fringes; TL: corolla tube length; TW: corolla tube width; ST: stigma exsertion; AN: anther exsertion; HT: display height; NF: number of flowers.

